# DeepSpot: a deep neural network for RNA spot enhancement in smFISH microscopy images

**DOI:** 10.1101/2021.11.25.469984

**Authors:** Emmanuel Bouilhol, Edgar Lefevre, Benjamin Dartigues, Robyn Brackin, Anca Flavia Savulescu, Macha Nikolski

## Abstract

Detection of RNA spots in single molecule FISH microscopy images remains a difficult task especially when applied to large volumes of data. The small size of RNA spots combined with high noise level of images often requires a manual adaptation of the spot detection thresholds for each image. In this work we introduce DeepSpot, a Deep Learning based tool specifically designed to enhance RNA spots which enables spot detection without need to resort to image per image parameter tuning. We show how our method can enable the downstream accurate detection of spots. The architecture of DeepSpot is inspired by small object detection approaches. It incorporates dilated convolutions into a module specifically designed for the Context Aggregation for Small Object (CASO) and uses Residual Convolutions to propagate this information along the network. This enables DeepSpot to enhance all RNA spots to the same intensity and thus circumvents the need for parameter tuning. We evaluated how easily spots can be detected in images enhanced by our method, by training DeepSpot on 20 simulated and 1 experimental datasets, and have shown that more than 97% accuracy is achieved. Moreover, comparison with alternative deep learning approaches for mRNA spot detection (deepBlink) indicated that DeepSpot allows more precise mRNA detection. In addition, we generated smFISH images from mouse fibroblasts in a wound healing assay to evaluate whether DeepSpot enhancement can enable seamless mRNA spot detection and thus streamline studies of localized mRNA expression in cells.

## 1 Introduction

High-resolution microscopy together with RNA single molecule fluorescence in-situ hybridization (smFISH) technologies allow gene expression profiling at subcellular precision for determining molecular states of various cell types (Ke et al., 2013) and that in high-throughput fashion (Battich et al., 2013). The repertoire of mRNA expression quantification methods is large and includes smFISH, clamp-FISH, amp-FISH and multiplexed versions such as e.g. MerFISH, all allowing the localization of RNA at sub-cellular level. There are technological differences between these methods in terms of the number of detected RNAs and number of processed cells, however all produce imaging data with mRNA spots that can be further matched to the spots’ x, y coordinates. With such increased image acquisition automation and the consequent growing number of high-throughput projects focused on spatially resolved transcriptomics, the need for automated and highly accurate detection of mRNA spots in fluorescent microscopy images has become increasingly important.

Despite the progress made in the recent years, accurately detecting the localization of spots corresponding to different mRNAs in a fully automatic fashion remains challenging. First, the background intensity is often irregular due to the fixative-induced fluorescence phenomenon (Rich et al., 2013); the signal quality is also dependent on the acquisition conditions and can produce motion blur or out-of-focus blur. Second, spot detection is affected by the non-homogeneous intensity distribution and indistinct spot boundaries relative to the background. Moreover, FISH images generally have a low signal to noise ratio (SNR), and the boundary between background (noise) and signal (spots) is not always obvious (Yano et al., 2017).

The main drawback of classical mRNA spot detection methods is the requirement of a strong human input to determine the best parameters to handle variable image to image properties such as SNR and presence of artifacts. Even small differences in these characteristics lead to the necessity for parameter fine-tuning (Caicedo et al., 2017). Other than being time-consuming, the quality of detection largely depends on the capacity of the user to correctly choose the method’s parameters according to each image properties (contrast, spots, artifacts, noise). Some recent deep-learning based approaches for mRNA spot detection try to circumvent this limitation, such as deepBlink (Eichenberger et al., 2021).

Here we introduce DeepSpot, a Convolutional Neural Network method dedicated to the enhancement of fluorescent spots in microscopy images and thus enabling downstream mRNA spot detection by conventional widely used tools without need for parameter fine-tuning. With DeepSpot we show that it is possible to avoid the manual parameter tuning steps by enhancing the signal of all spots so that they have the same intensity throughout all images regardless to the contrast, noise or spots shape.

DeepSpot gives a new twist to the ResNet network architecture and learns to automatically enhance the mRNA spots, bringing them all to the same intensity. In parallel, a multi-network architecture is integrated, trained by minimizing the binary cross-entropy (BCE) while providing context for mRNA spots thanks to the atrous convolutions. We evaluated the impact of the spot enhancement on the downstream mRNA spot detection, by performing spot detection using ICY with fixed parameters on both simulated images and experimental images manually annotated. Moreover, we compared the quality of mRNA spot detection from images enhanced by DeepSpot with deepBlink, and have shown that our method achieves greater generalization to effectively handle full variability of smFISH data. Finally, to illustrate the end-to-end use of DeepSpot in projects where detecting sub-cellular localization with high precision is essential, we generated smFISH images from mouse fibroblasts in a wound healing assay, where enrichment of expression of *β-Actin* towards the location of the wound is expected in the migrating 3T3 mouse fibroblasts.

## 2 Related Work

Methodologically mRNA spot detection can be positioned at the intersection of two related topics in image analysis: detection of small blobs and detection of small objects. The goal of blob detection is to find regions in an image that differ from the surroundings with respect to certain properties, such as brightness or shape. More precisely, a blob is a region with at least one local extremum (Lindeberg, 1993). Blobs can be considered as a particular case of more or less circular objects. Object detection has been one of key topics in computer vision which goal is to find the bounding box of objects. However, small object detection, such as mRNA spots, remains difficult because of low-resolution and limited pixels (Huang et al., 2017).

In this work we propose a deep learning network inspired by small object detection approaches for mRNA spot enhancement and we show how it can enable the downstream accurate detection of spots.

### 2.1 mRNA spot detection

The specificity of mRNA spot detection is that the corresponding blobs are necessarily small. Ideally, mRNA spots would correspond to the maximum intensity pixel surrounded by the diffraction of the fluorescent signal that can be modeled by a Gaussian within a disk of small radius (see Figure 1, image B). This radius is dependent on multiple acquisition parameters and optical properties of the microscope such as the diffraction limit, the fluorophore excitation state and the Point Spread Function (PSF). Typically, in the case of FISH images, mRNA spots are small, compact and below the resolution limit of the microscope (Olivo-Marin, 2002).

**Figure 1:**
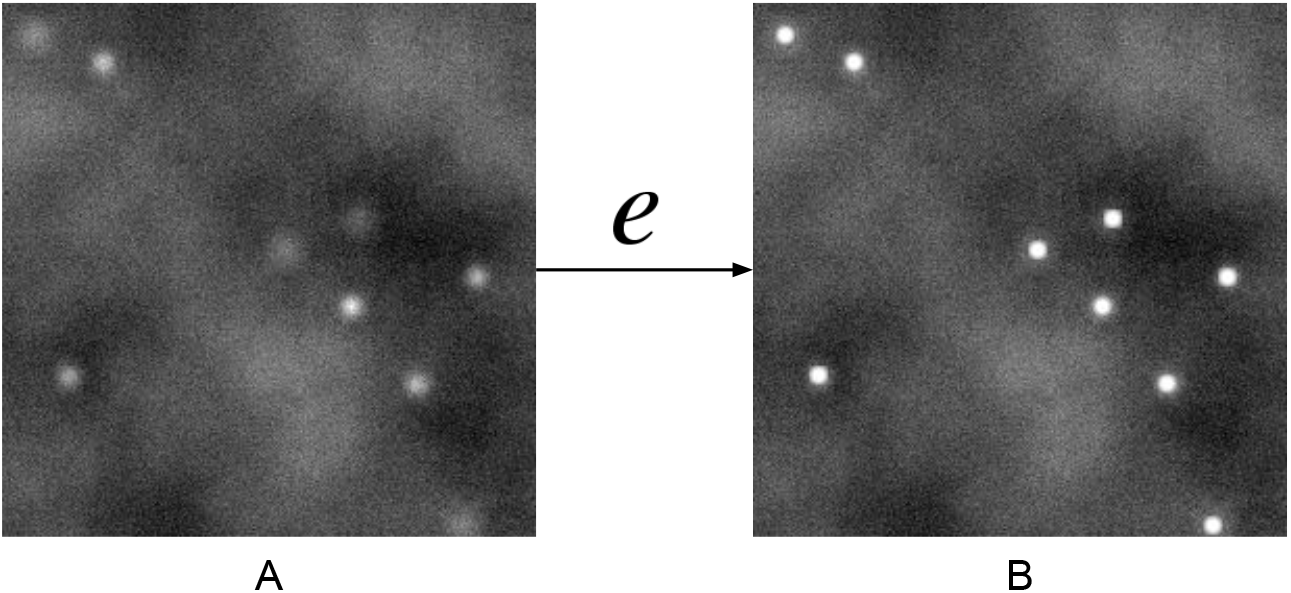
RNA spots on a noisy background (image A). Spots’ intensity is increased after the enhancement by *e* (image B).

While there is no universal solution to the detection of small blob-like objects such as mRNA spots in fluorescent cellular images, a large plethora of work is available on the subject. A number of approaches have gained wide popularity thanks to the development of software tools embedding the algorithms and providing user with a graphical interface. In particular, ImageJ/Fiji (Abràmoff et al., 2004; Schindelin et al., 2012) is widely used, largely due to the plugin-based architecture, recordable macro language and programmable Java API. Another popular tool is CellProfiler (Kamentsky et al., 2011), based on similar paradigms and FISH-Quant (Mueller et al., 2013). A more recent platform, ICY (De Chaumont et al., 2012), also provides the possibility to develop new algorithms as well as a user interface for image analysis, including the ICY spot detector for the detection of mRNA spots, method based on wavelet transform decomposition (Olivo-Marin, 2002).

Deep-learning networks have been introduced for mRNA spot detection (Gudla et al., 2017; Mabaso et al., 2018), and more recently deepBlink (Eichenberger et al., 2021). The latter focuses on a fully convolutional neural network based on the U-Net architecture. deepBlink not only provides the code, but also annotated smFISH data and implements a threshold-independent localization of spots.

### 2.2 Detection of small objects

Consistently with the mRNA spot detection difficulty, detection of small objects remains a challenging part of the general object detection problem due to the limited information contained in small regions of interest. For instance, it has been shown that the object size has a major impact on the accuracy of Deep Learning object detection networks such as VGG, ResNet or Inception V2 (Huang et al., 2017). Indeed, small objects do not contain sufficient semantic information (Liu et al., 2021) and thus the challenge is to capture semantic features while minimizing spatial information attenuation (Fu et al., 2020).

Expectedly, adding more context improves the detection of small objects (Bell et al., 2016; Fu et al., 2020; Noh et al., 2019). An elegant solution is to use the dilated convolution (a.k.a. atrous convolution), because the receptive field can be expanded without loss of resolution and thus capture additional context without loss of spatial information (Hamaguchi et al., 2017; Yu and Koltun, 2015).

### 2.3 Small blob enhancement

Of particular interest to our work is the signal enhancement, an image processing technique aiming to reinforce the signal only in those regions of the image where the actual objects of interest are, and potentially to weaken the noise or the signal from other structures (Smal et al., 2009). In our case, the objects of interest are small blobs corresponding to mRNA spots.

Image enhancement is the transformation of one image *X* into another image *e*(*X*) (see Figure 1). Pixel values (intensities) of *X* at spot locations are modified according to the transformation function *e*, the resulting pixel values in image *e*(*X*) measuring the certainty of mRNA presence at that position. Thus, *e*(*X*) can be considered as a probability map that describes possible mRNA spot locations (Smal et al., 2009).

Small blob enhancement has been developed in other fields than mRNA spots in fluorescent imaging, such as in astronomy to enhance stars or galaxies over the cosmic microwave background (Sadr et al., 2019) or in the biomedical image to facilitate the human detection of small blob structures such as nodules (Liu et al., 2010). However, these images don’t have the same characteristics as smFISH data in terms of noise or signal. For example, human nodules are much larger objects than typical mRNA spots. As for the star enhancement method proposed by (Sadr et al., 2019), it is not suited for low intensities blobs (Pino et al., 2021), which is a major concern in spot detection for smFISH images.

## 3 Materials and Methods

### 3.1 Materials

#### 3.1.1 smFISH data

We constructed a dataset DS_exp_ of 1553 images from the experimental smFISH data acquired in (Chouaib et al., 2020), by applying 256 × 256 pixel patches to better fit in the GPU memory (see Table 1). The authors have performed spot detection using image analysis techniques such as local maximum detection, implemented in the BIG-FISH pipeline (Imbert et al., 2021). Along with the scripts, the authors provided a list of 57 combinations of parameters to detect the spots in their study. We run the pipeline with these parameters and performed an additional manual curation to keep patches with number of spots between 10 and 150 and remove those with a visually obvious over- or under-identification of spots. The resulting DS_exp_ dataset of 1553 images is thus experimentally generated and guarantees high confidence in the ground truth annotation of mRNA spot coordinates.

**Table 1:** List of datasets used for training (22 training datasets) and evaluation (21 test datasets) of the DeepSpot network. Images in the 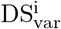 datasets have spot intensities between 80 and 150 for each image. For the fixed intensity datasets 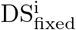 the spot intensity is set to one value within [80..150] for a given image, but varies from image to image. 10 variable intensity 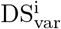 and 10 fixed intensity 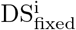 datasets are named according to the number of spots in the images, *i*. The dataset DS_hybrid_ combines DS_exp_ with 25% simulated images.

| Datasets | #spots<br>per image | intensity | #images<br>train | #images<br>test | #datasets<br>train/test |
| --- | --- | --- | --- | --- | --- |
| DS<sub>exp</sub> | $10 \leq i \leq 150$ | variable | 1453 | 100 | 1/1 |
| DS<sub>var</sub><sup>i</sup> | $i \in \{[10..100] \bmod 10\}$ | variable | 2000 | 400 | 10/10 |
| DS<sub>fixed</sub><sup>i</sup> | $i \in \{[10..100] \bmod 10\}$ | fixed | 2000 | 400 | 10/10 |
| DS<sub>hybrid</sub> | $10 \leq i \leq 150$ | variable | 2000 | NA | 1/NA |

To evaluate whether DeepSpot enables precise mRNA spot detection in a biological context, we made use of the wound healing assay (Moutasim et al., 2011) to generate a second experimental dataset DS_wound_. In the wound healing assay, migrating cells, such as fibroblasts are grown on a coverslip and serum starved for synchronization. A scratch in the middle of the coverslip is then generated, mimicking a wound, followed by induction of the cells to polarize and migrate towards the wound, to generate wound closure, done using replacement of serum starved medium with 10 % FBS-containing medium. We used 3T3 mouse fibroblasts in a wound healing assay, followed by cell fixation. Fixed samples were taken for single molecule FISH experiments to visualize and quantify *β-Actin* mRNA and imaged on a custom built Nikon Ti Eclipse widefield TIRF microscope. *β-Actin* has been previously shown to be enriched in neuronal growth cones of extending axons, as well as the leading edges of migrating cells and this enrichment has typically been associated with cell polarity and neuronal plasticity (Zhang et al., 1999; Lapidus et al., 2007; Condeelis and Singer, 2005). Based on this, we hypothesized that *β-Actin* would be enriched in the leading edge of migrating 3T3 fibroblasts. The dataset is composed of 96 images, 48 images of non migrating cells (control) and 48 images of migrating cells. Each image was divided into 4 patches, yielding a total of 384 patches of 256 × 256 pixels size.

#### 3.1.2 Simulated, experimental and hybrid dataset

In addition to the experimental datasets, we have built 20 simulated datasets with images 256×256 pixels of width and height, same as the patches of DS_exp_. Briefly, the background was generated by a combination of Poisson noise and Perlin noise to which was added a motion blur modeled by elastic transformations applied to the noise. Spots were generated as circles, their size between 8 and 15 pixels in diameter. They were randomly placed in the image and then convolved by a Gaussian function approximating PSF.

Two different types of datasets were generated DS_fixed_ and DS_var_, each containing 10 datasets defined according to the number of spots per image *i* ∈ {[10..100] mod 10}, see Table 1. For example, in the 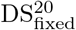 dataset each image contains 20 spots and in the 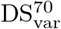 dataset each image contains 70 spots. Each image in the DS_fixed_ dataset has the same fixed spot intensity for all spots randomly chosen in the interval [80..150], while in the DS_var_ dataset the spot intensity is randomly chosen from the same interval for each spot, resulting in images with variable spot intensity.

In addition to the experimental and simulated datasets, we have built a hybrid dataset DS_hybrid_ where the experimental data from DS_exp_ is augmented by appending an additional 25% of simulated images generated with both variable spots’ intensity and variable spots’ number per image, within the [80, 150] and [10, 100] intervals, respectively.

#### 3.1.3 Ground truth

Since the goal of our network is to learn to transform an image *X* into *e*(*X*), where intensity at spots’ location is enhanced, the training step has to be provided with the enhanced counterpart of each image in the training set. That is, training sets include pairs of images ⟨*X, e*(*X*)⟩ where *e* is the procedure that is used to produce ground truth enhanced images: for each spot (Figure 1, image A), the ground truth enhancement procedure *e* is applied at the spots’ locations *A*(*X*), resulting in images where spots are enhanced as shown in Figure 1, image B.

In this work we implement *e* as a kernel of 3 × 3 pixels at all locations where the spots were annotated in the experimental dataset DS_exp_ or generated for DS_fixed_, DS_var_ and DS_hybrid_ (see Table 1. The kernel has the same pixel values for all the spots, in order to drive the network to learn to enhance all spots up to the same level of intensity, regardless of the initial intensity in the acquired data.

### 3.2 Method

In this section, we present the DeepSpot enhancement network in detail. We first overview the network architecture, and then we discuss the custom loss function.

#### 3.2.1 Network architecture

DeepSpot network is composed of two main components as presented in Figure 2. The first component is a multi-path network shown in panel A. The second component is an adapted residual network as shown in panel B.

**Figure 2:**
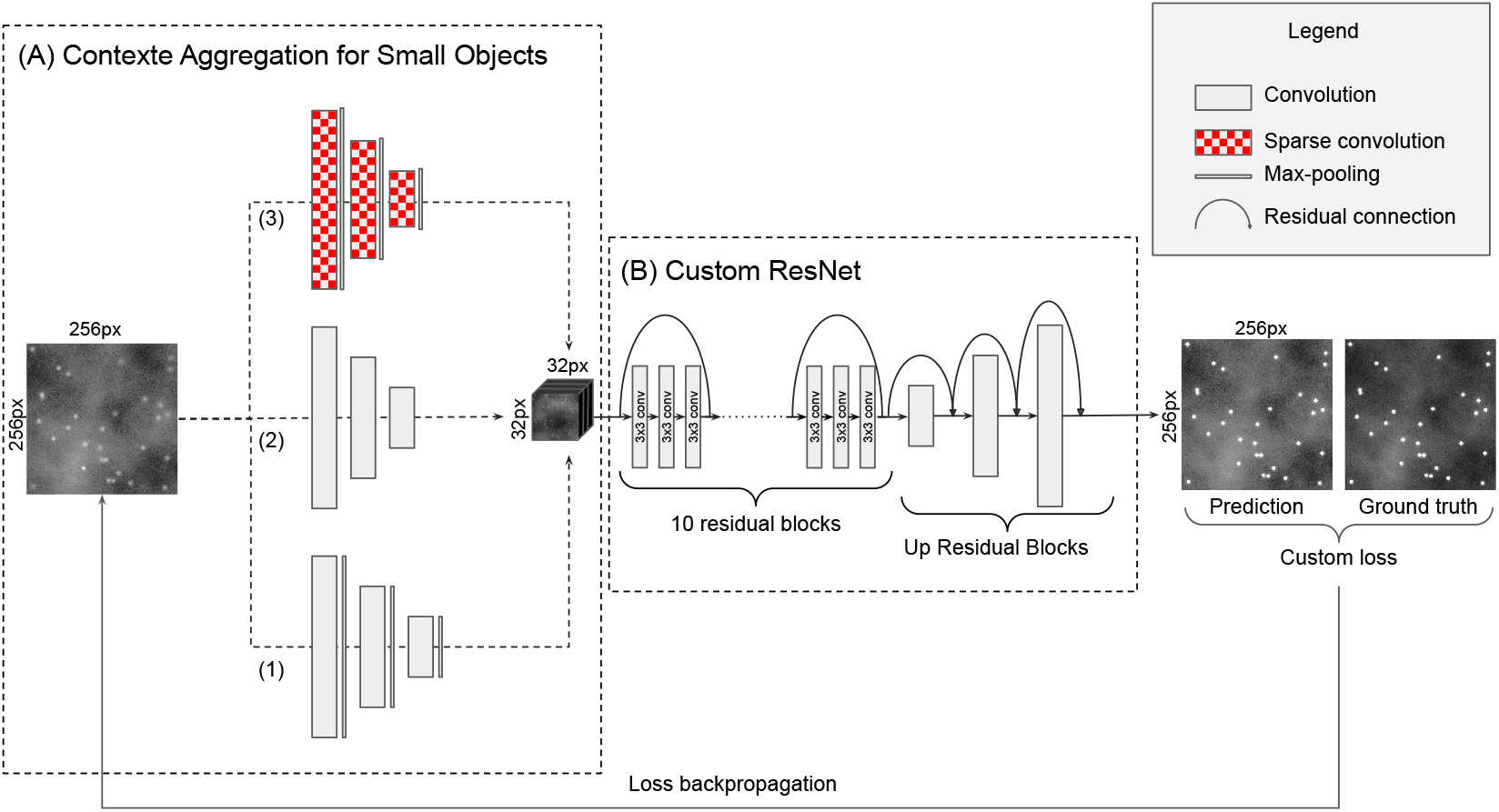
DeepSpot network architecture is composed of the Context Aggregation for Small Objects module (CASO) constituted of a multi-path network (panel A) and a customized ResNet component (panel B). A custom loss function is used for training the network.

##### Context Aggregation Module

As pointed out in (Hamaguchi et al., 2017), finding small objects is fundamentally more challenging than large objects, because the signal is necessarily weaker. To solve this problem in the contexte of mRNA spot detection, we developed a new module, that we called the Context Aggregation for Small Objects module (CASO). (Hu and Ramanan, 2017) demonstrated that using image evidence beyond the object extent (context) always enhances small object detection results, we therefore developed our CASO module to aggregate context around the mRNA spots. The CASO module is a multi-path network as shown in panel A. It takes the input image and processes it along three different paths each with different types of convolution blocks to collect specific information from the input image. Each path contains 3 convolution blocks.

1. The first path is composed of traditional convolution blocks (2D convolution, batch normalization, activation and max pooling). These blocks, often used in CNNs, are particularly efficient to reinforce the semantic information at the expense of spatial information.
2. The second path uses only 2D convolutions, batch normalization and activation. As Max Pooling is known to keep mostly the maximum intensities in images, some of the faint spots may be eliminated during the max pooling operation. In this path we used strided 2D convolutions instead of a max pooling layer, to keep the information of low intensity of spots. To make sure to end up with the same receptive field as the first path, we set the stride to 2.
3. The third path makes use of the atrous convolution pooling (Hamaguchi et al., 2017), implemented as a 2D convolution with the dilatation rate of 2. The following layers are batch normalization, activation and max pooling.

The CASO module is a multi-path neural network and can learn more comprehensive and complementary features than a single path. In particular, the goal of the atrous convolution is to bring more context around the small spots (see section 2.2), while the two other paths aggregate the semantic information of the bright spots and faint spots for the first and second path respectively. The results of the three encoding paths are then concatenated to construct a longer feature vector containing information extracted by each path. For all convolutional blocks, the activation function is the Rectified Linear Unit (ReLU). The number of filters for the 2D convolutions in the are 32, 64, 128 for the first, second and third block, respectively.

##### Custom ResNet

For the second component (panel B) we customized the ResNet architecture to create a residual neural network composed of ten consecutive convolutional residual blocks (ResBlock), using full pre-activation blocks described in (He et al., 2016), where the authors suggested that the better results obtained by the full pre-activation blocks are due to the pre-activation by the batch normalization that improves regularization of the model due to the fact that the inputs to all weight layers have been normalized. Each ResBlock is composed of three sub-blocks as presented in Figure 3. A sub-block is constituted of a batch normalization, followed by an activation (ReLU) and a 2D convolution. After the 3 sub-blocks a spatial dropout layer with rate 0.2 is applied. Each ResBlock ends by the residual connection.

**Figure 3:**
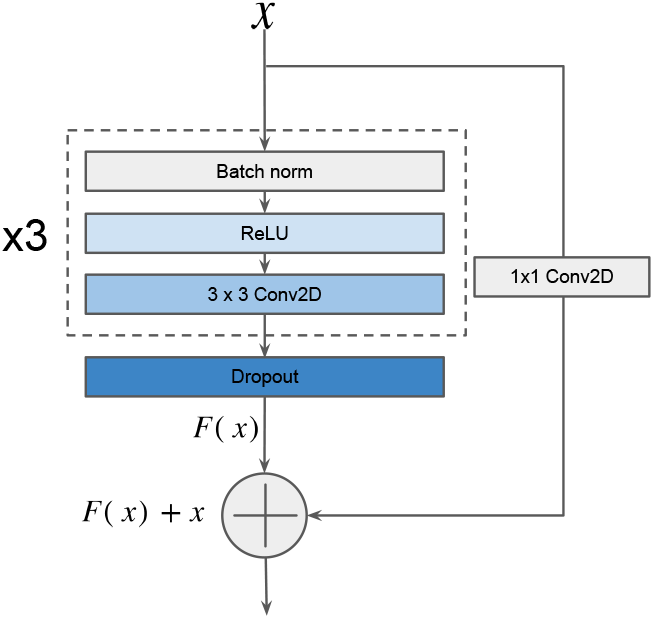
Full pre-activation residual block, composed of Batch normalisation, Activation, Convolution, repeated three time before Dropout and residual connection.

To obtain an output image with the same size as the input image, we used a particular type of up convolutional blocks. Recently, (Wang et al., 2021) demonstrated that the use of Up Residual Blocks (UpResBlocks) instead of classic up-convolution blocks improves the performance of generative networks by preserving the effective features from the low dimensional feature space to the high dimensional feature space. Our decoding path is composed of three UpResBlocks and reconstitutes an output with the same size as the input while propagating low dimension feature information of the enhanced spots from the custom ResNet to the last layer. Each UpResBlock is constituted of three sub-blocks containing a 2D transposed convolution, batch normalization and activation (ReLU). The three sub-blocks are followed by a spatial dropout layer with rate 0.2. UpResBlocks then end with a residual connection. A sigmoid activation function is applied to the last convolution, so that all the pixels have values in the [0, 1] interval. The final image is obtained by normalizing the pixel intensities between 0 and 255.

#### 3.2.2 Loss

We defined our custom loss function as a combination of binary cross-entropy (BCE) and Mean Squared Error (MSE) functions. The main term of the loss function is the BCE loss, defined by 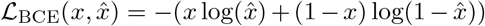 that measures the difference between the images predicted by the network 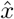 and the ground truth images *x*. While mostly used for classification, it can also be used for segmentation and enhancement due to its performance for pixel-level classification (Jadon, 2020).

To this main ℒ_BCE_ term we added a regularization term defined by Mean Squared Error, 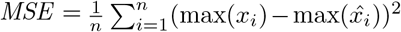 that is computed between the maximum value of the predicted image 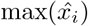 and the maximum value of the ground truth image max(*x*_*i*_). This regularization drives the network to produce spots whose intensity is close to 255 (see Table 2), and therefore standardizes the signal enhancement intensity in the output images, which in its turn facilitates the downstream automatic detection of the spots. The total loss function is ℒ_BCE_ + ℒ_MSE_.

**Table 2:** Spot enhancement performance in terms of resulting spot intensity. The measures displayed correspond to the spot intensity between [0,255] after enhancement by the neural network and averaged by category of models and datasets. Between brackets are shown the 95% confidence intervals. In rows are listed the categories of models, columns correspond to the dataset categories on which the different models were applied.

| Train \ Test | DS<sub>exp</sub> | DS<sub>fixed</sub> | DS<sub>var</sub> |
| --- | --- | --- | --- |
| M<sub>exp</sub> | 249.55<br>[249.29, 249.82] | 250.81<br>[250.68, 250.93] | 247.07<br>[246.68, 247.45] |
| M<sub>hybrid</sub> | 241.15<br>[240.79, 241.52] | 248.71<br>[248.57, 248.85] | 243.3<br>[243.14, 243.45] |
| M<sub>var</sub> | 248.92<br>[248.47, 249.37] | 251.99<br>[251.93, 252.05] | 254.57<br>[254.53, 254.62] |
| M<sub>fixed</sub> | 242.93<br>[242.16, 243.71] | 254.73<br>[254.71, 254.75] | 249.33<br>[249.09, 249.57] |

## 4 Results

We trained the DeepSpot network on our 20 simulated training datasets DS_fixed_ and DS_var_ as well as on the experimental and hybrid datasets DS_exp_ and DS_hybrid_, resulting in 22 models M_fixed_, M_var_, M_exp_ and M_hybrid_. Training parameters were optimized with HyperOpt algorithm (Bergstra et al., 2013) and the ASHA scheduler (Li et al., 2020). The best configuration obtained and used for further trainings had the learning rate of 0.0001, the dropout rate of 0.2 and the batch size of 32 and 128 filters per convolution.

Each of the resulting 22 models was evaluated on the 21 test data sets from Table 1, yielding 462 enhanced test datasets (8100 images total). To assess whether DeepSpot enhancement enables easy spot detection, we applied the ICY spot detector (Olivo-Marin, 2002) to the images enhanced by different models. Moreover, we defined an unique set of ICY parameters that matches the shape and intensity of the enhancing kernel of the DeepSpot network: scale 3, sensitivity 20 and scale 7, sensitivity 100. We then evaluated whether the detected spots from the enhanced images matched well with the annotated ground truth of spots’ coordinates.

### 4.1 Evaluation procedure

We denote by *D*(*X*) = {*p*_1_, … *p*_*n*_} the point pattern detected by ICY from the enhancement of of an image *X* by DeepSpot and by *A*(*X*) = {*q*_1_, … *q*_*m*_} the ground truth annotation of spots coordinates. Notice that *m* is not necessarily equal to *n*, corresponding to under- or over-detection and even for a well-detected spot, the coordinates in *A* and *D* may slightly differ. To account for these remarks, we used the *k*-d tree algorithm (Bentley, 1975) to query the detection *D* for nearest neighbors in *A* as proposed in (Samet, 1990; De Berg et al., 2008). Number of neighbors was set to 1 and matching radius *t* to 3 (coherent with the enhancement kernel for ground truth images).

This allows to establish a matching for all annotated points under *t* = 3 and thus also defines the number of False Negatives or False Positives corresponding to the missing matches from *D*(*X*) or *A*(*X*), respectively. True Negatives are defined by all pixels *p* of the confusion matrix such that *p* ∈ *X* \ {*A*(*X*) ⋃ *D*(*X*)}. However, given that |*X*| ≫ |*X* \ {*A*(*X*) ⋃ *D*(*X*)}| implies inflated TN values, this makes measures such as accuracy, AUC and ROC curve irrelevant.

The potential drawback of the *k*-d tree approach is that two points *q*_*i*_, *q*_*j*_ ∈ *A* can match to one *p*_*k*_ ∈ *D* (see Figure 4). This can happen if annotated spots *q*_*i*_, *q*_*j*_ are close and the detected matching point for both of them, *p*_*k*_, lies within the same distance *t*, which can correspond to an over enhancement and thus blurring between the two spots in the enhanced image. To measure this effect we thus also report the number of ambiguous matches (AM).

**Figure 4:**
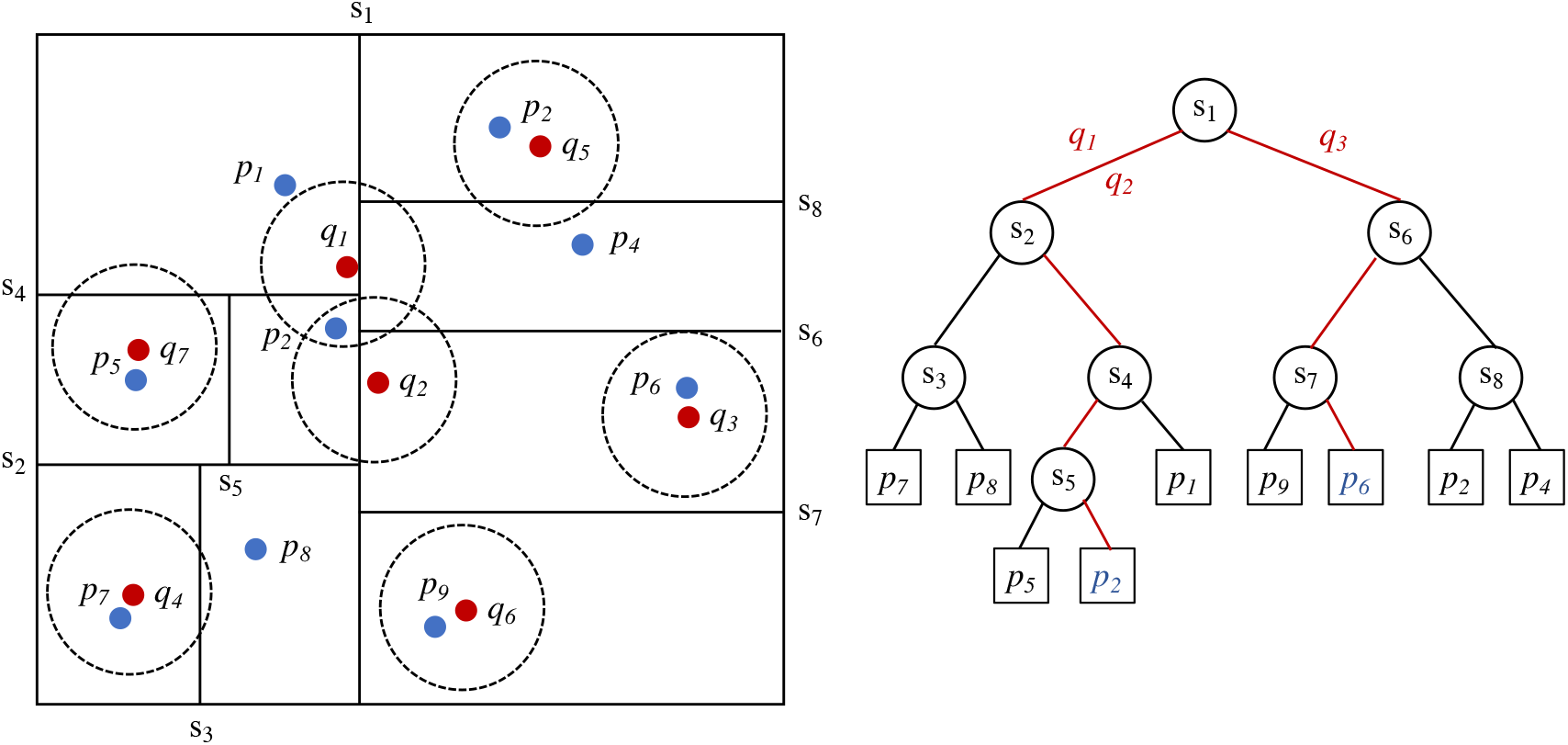
Spot matching by 1-neighbor *k*-d tree between the detected mRNA spots {*p*_1_, …, *p*_9_} depicted in blue and annotated spots {*q*_1_, …, *q*_7_} depicted in red. The *k*-d tree construction for {*p*_1_, … *p*_9_} is shown on the left. Using the matching radius depicted by circles, the *k*-d tree queries for *q*_1_ and *q*_2_ shown in red, lead to the same leaf *p*_2_ and correspond to an ambiguous match, while query for *q*_3_ leads to an unique match. mRNA spots *p*_1_, *p*_4_ and *p*_8_ are the False Negatives.

### 4.2 DeepSpot enhances the mRNA spot signal to the same intensity

To avoid the manual selection of the detection threshold, it is imperative to have a homogeneous spot intensity for the whole dataset in order to use unique set of parameters for all images. Table 2 summarizes the intensities obtained after enhancement by DeepSpot for each category of datasets described in Table 1 (experimental, hybrid, simulated with variable and fixed intensities).

As expected, intensities were closer to 255 when training and test sets data belong to the same data category. For example, models trained on data with fixed intensities M_fixed_ and applied to data with fixed intensities DS_fixed_ produced enhanced spot intensities very close to 255. Similarly enhancement close to 255 could be observed when models M_var_ were evaluated on DS_var_ Of particular interest are the enhancement results from the M_exp_ training, that are close to 250 for every dataset, experimental or simulated. Hybrid model enhances intensities to 241 on the experimental dataset. In general the enhanced spot intensities were between 241 and 255, representing a variation of only 5.4% from the maximum intensity which is sufficient to fully separate smFISH spots from the background in the enhanced images.

### 4.3 DeepSpot enables accurate mRNA spot detection

Summary statistics of performance of each model type are shown in Table 3 including the mean F1-score, precision, recall and ambiguous matches, with 95% confidence interval. For a given model *M* each metric was computed for all enhanced images (8100 total). The mean metric value *x* and 95% confidence interval *C* were then calculated separately for each model type. Due to the high prevalence of True Negatives, instead of the Accuracy measure, we calculated the F1-Score, which gives an indication of the model accuracy with a better balance between classes than the actual accuracy measure, by not including True Negatives.

**Table 3:** Models’ performance per model type. Metrics (F1-score, precision, recall, Ambiguous Matches (AM)) were calculated by averaging the values obtained for each image of the 21 test datasets. Top values in cells correspond to the mean value, bottom values between brackets show the 95% confidence interval. Best values are highlighted in bold.

| Model | F1-Score(%) | Precision(%) | Recall(%) | AM(mean/img) |
| --- | --- | --- | --- | --- |
| $M_{\text{exp}}$ | 87.8<br>[87.6, 88.1] | 99.6<br>[99.5, 99.7] | 80.1<br>[79.8, 80.5] | 2.7<br>[1.4, 4.0] |
| $M_{\text{hybrid}}$ | **97.3**<br>[97.2, 97.3] | 99.3<br>[99.2, 99.4] | **95.5**<br>[95.4, 95.6] | **0.94**<br>[0.56, 1.32] |
| $M_{\text{var}}$ | 94.3<br>[94.3, 94.4] | 99.5<br>[99.5, 99.5] | 91.4<br>[91.3, 91.5] | 1.19<br>[1.03, 1.35] |
| $M_{\text{fixed}}$ | 91.6<br>[91.5, 91.7] | **99.9**<br>[99.9, 99.9] | 85.7<br>[85.6, 85.8] | 2.05<br>[1.77, 2.32] |

The results in Table 3 indicate that the number of FP is very low for all models, given that both precision and recall are high. Importantly, M_hybrid_ has shown best best overall performance in terms of precision, recall and F1-score, thus indicating that the mRNA spot enhancement by M_hybrid_ leads to the least FP and FN counts in the downstream mRNA spot detection.

Figure 5 shows the mean F1-scores for each of the 22 models evaluated on the 21 datasets. This heatmap indicates that (i) models trained on the 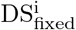 training sets perform better on the corresponding 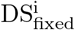 test sets rather than on variable intensity test sets, (ii) models trained on 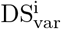 training sets show better performance on the corresponding test sets rather than on fixed intensity test sets. It also shows that M_var_ models are globally better than M_fixed_ models. A plausible hypothesis is that training on variable intensities makes models better at generalizing on other data. Finally, M_hybrid_ is the model that has the best overall performance, including the experimental dataset. Again, the diversity of training data drives this model to be more robust to newly encountered data.

**Figure 5:**
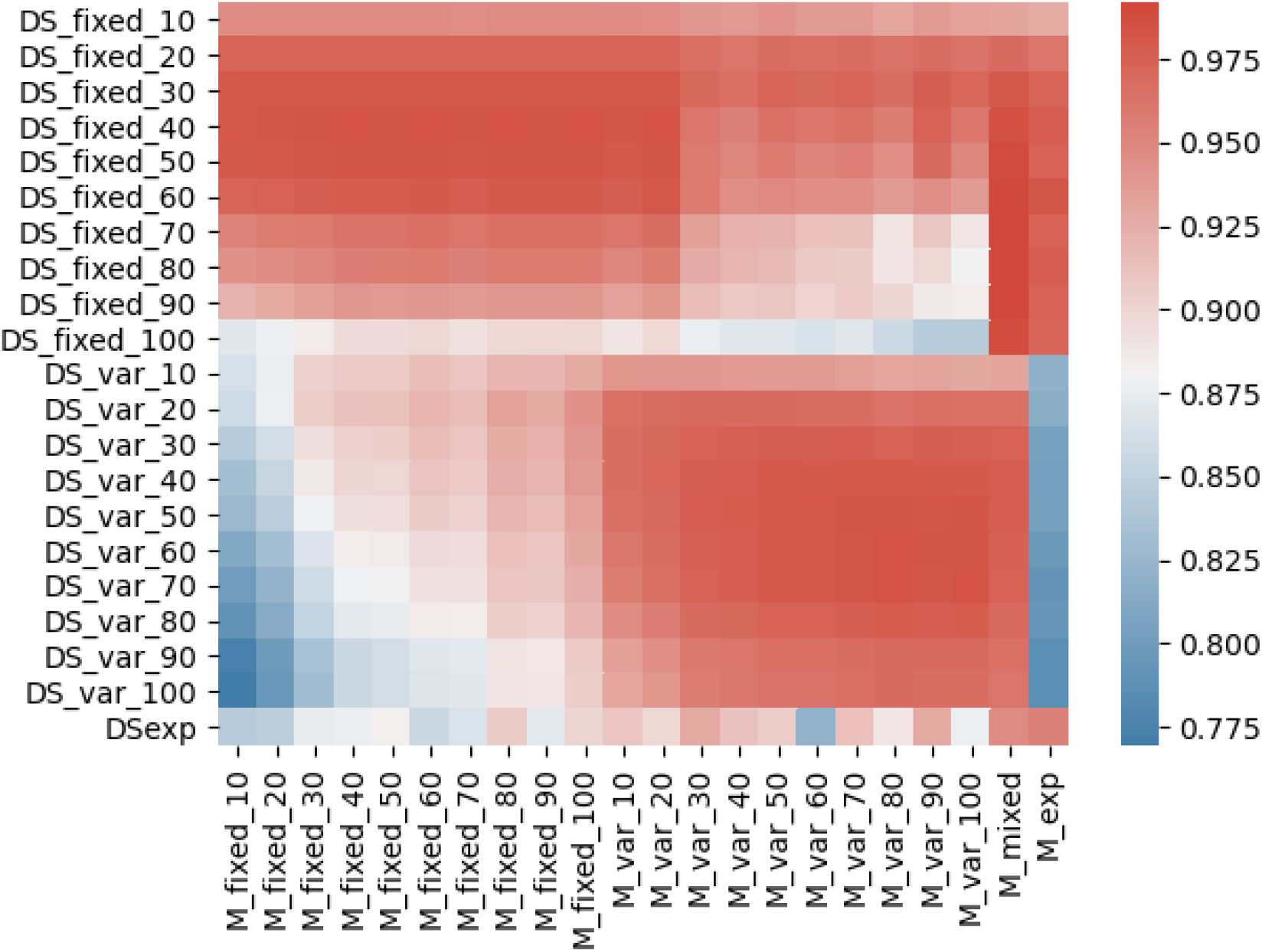
Heatmap of the F1-scores obtained by each of the 22 models when evaluated on the 21 test datasets described in Table 1

More generally, given that the F1-score is above 90% for all the models, we can conclude that the architecture of the DeepSpot neural network is particularly suited for the task of mRNA spot enhancement.

### 4.4 DeepSpot enables more accurate spot detection compared with deepBlink

The main objective of our DeepSpot method is to circumvent the parameter fine-tuning and enable the downstream spot detection with an unique parameter set. As such, this objective fits well with the one that the authors of deepBlik (Eichenberger et al., 2021) have pursued, despite the fact that the latter proposes a new spot detection method, while our goal is to fit a spot enhancement step into commonly used workflows. Consequently, deepBlink constitutes a relevant comparison target.

We compared the accuracy of spot detection by the model M_dB_ made available on deepBlink associated GitHub with that of DeepSpot when trained on hybrid data M_hybrid_, both on our datasets as well as on the dataset provided by the authors of deepBlink, DS_dB_. Table 4 shows the F1-scores for each dataset category.

**Table 4:** Models’ performance for deepBlink and DeepSpot for smFISH spot detection. Overall F1-scores are calculated by averaging the values obtained for each image of the test datasets corresponding to each dataset category. Top values in each cell correspond to the mean value, bottom values between brackets show the 95% confidence interval. Best values are highlighted in bold.

| Model \ F1-score(%) | $DS_{exp}$ | $DS_{var}$ | $DS_{fixed}$ | $DS_{dB}$ |
| --- | --- | --- | --- | --- |
| $M_{dB}$ | 78.7<br>[75.26, 82.14] | 47.97<br>[46.86, 49.07] | 67.4<br>[66.28, 68.51] | **94.2**<br>+- 0.141(std) |
| $M_{hybrid}$ | **94.43**<br>[93.41, 95.45] | **96.67**<br>[96.58, 96.75] | **98.04**<br>[97.96, 98.12] | 87.92<br>[85.36, 90.49] |

DeepSpot clearly outperformed deepBlink on our datasets (Table 4). However, we found that M_dB_ performed better on experimental data than on simulated data, presumably because deepBlink model has only been trained on experimental data. M_hybrid_ results were consistent on all of our datasets. To complete the comparison, we applied our M_hybrid_ model to the smFISH test dataset DS_dB_ made available with the deepBlink publication and containing 129 smFISH images. The results reported for the prediction of M_dB_ for the DS_dB_ dataset is the result obtained by the authors and reported in (Eichenberger et al., 2021). Not surprisingly, deepBlink model performance is better on their own smFISH images, however DeepSpot managed to have an F1-score of nearly 88%, a noticeable achievement since the model has not been trained on the DS_dB_ data. Together, the results of table 4 indicated that DeepSpot is a robust methodology that offers a generalist model for mRNA spot enhancement and ensures high quality downstream spot detection without parameter tuning.

### 4.5 DeepSpot’s use in an end-to-end smFISH experiment

To evaluate whether DeepSpot can be effectively used in an and-to-end smFISH experiment, we have performed a wound healing assay in which cells migrate towards a wound to close it. To investigate whether *β-Actin* was enriched at the leading edges of 3T3 migrating fibroblast cells (see 3.1.1). The wound location was manually annotated as shown on a typical image example in panel A, Figure 6.

**Figure 6:**
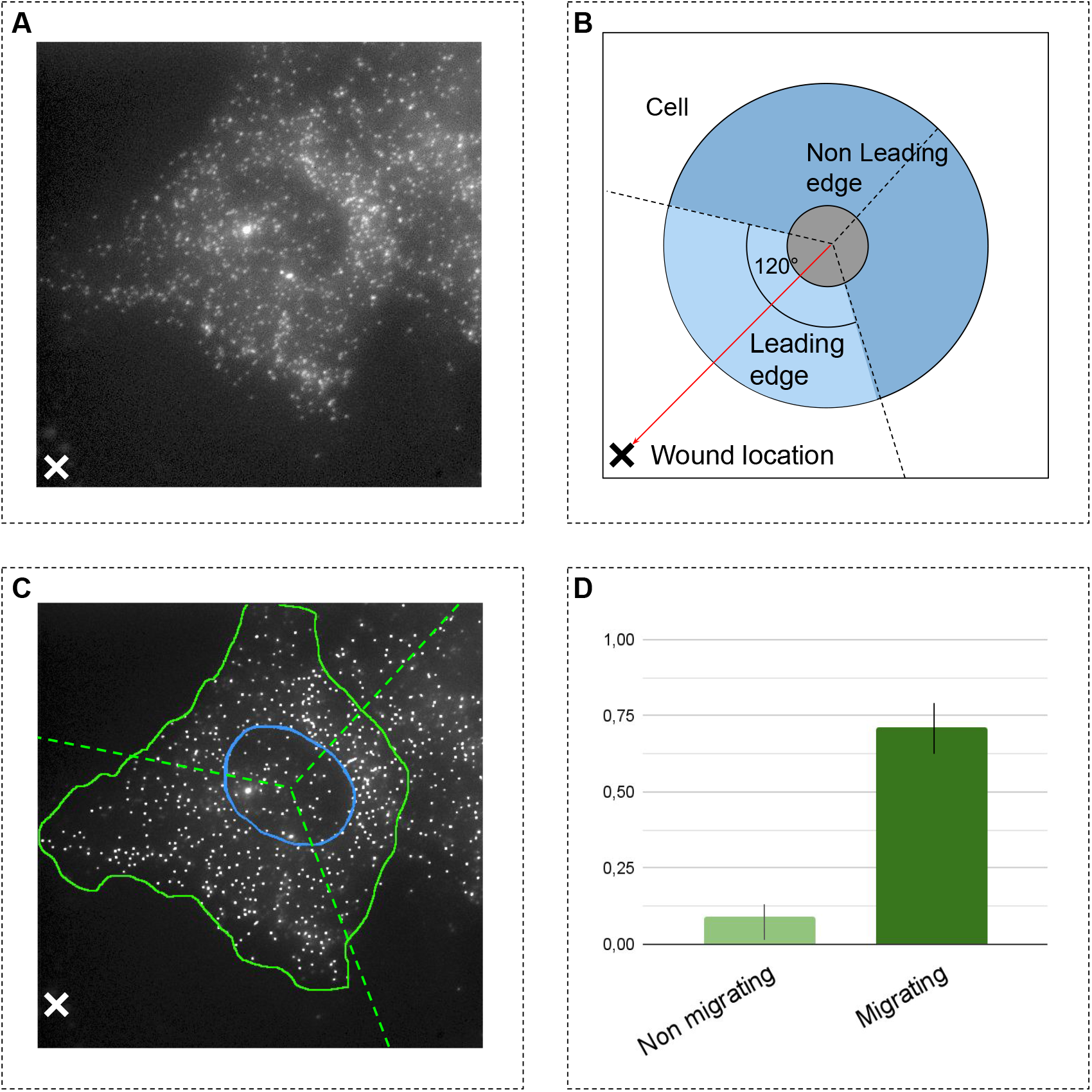
Processing steps for the end-to-end wound healing assay. Panel A is a typical smFISH image of the wound healing assay is shown. Panel B shows the cell quantization procedure in 3 sections and the direction of the wound (red arrow). S_wound_ section is oriented towards the wound and is shown in light blue, other sections are in dark blue. Panel C represent how th cell and nucleus were manually segmented and mRNA spots were counted after enhancement within the cytoplasmic portion of each section. Panel D presents the Wound Polarity Index (WPI) of cytoplasmic mRNA transcripts in S_wound_ compared to other sections for the *β-Actin* RNA. WPI was calculated in migrating and non-migrating cells.

All the cell images were segmented by manually, yielding cell and nucleus masks. Wound location defined the cell migration direction as schematically shown in panel C, Figure 6. We partitioned the cell masks into three section by computing 120° section centered at the nucleus centroid, and anchoring one of these section as oriented towards the wound location S_wound_ at 60° angle to the left and to the right of the line between the nucleus centroid and the wound location. This cell segmentation allowed us to compute the normalised number of detected mRNA spots in the cytoplasmic part of each section *S*_1_, *S*_2_, *S*_3_.

Using the DypFISH framework (Savulescu et al., 2021), we further compared the cytoplasmic mRNA relative density in S_wound_ (light blue) and in sections that are not oriented towards the wound (dark blue), as shown in Figure 5, Panel B. We used the Polarity Index (Savulescu et al., 2021), that measures the enrichment of mRNA in different sections. Briefly, the Polarity Index measures how frequently the relative concentration within the wound section is higher than in the non-wound section. The Polarity Index lies between [−1, 1], a positive value implying a wound-correlated enrichment of RNA transcripts while a negative values implies enrichment away from the wound and a value of zero implies no detectable enrichment.

*β-Actin* mRNA was highly enriched in the leading edge of migrating cells, whereas almost no detectable enrichment of *β-Actin* was found in the leading edge of control cells. This is in line with previously published data, showing enrichment of *β-Actin* in leading edges of migrating fibroblasts (Kislauskis et al., 1993).

## 5 Discussion and conclusion

Recent FISH microscopy methods are capable of generating thousands of images, it has thus become imperative to introduce algorithms capable to streamline the detection of mRNA spots and in particular to avoid manual fine-tuning of numerous parameters.

In this work we introduced DeepSpot, a novel CNN architecture specifically designed to enhance RNA spots in FISH images, thus enabling the downstream use of well known spot detection algorithms, such as the ICY spot detector, without parameter tuning. In particular, the architecture of our network introduces the Context Aggregation for Small Object module that relies on sparse convolution to provide more context for enhancement of small blob-like objects corresponding to mRNA spots. DeepSpot network has been trained and tested on 21 simulated datasets, all with different signal and noise characteristics, as well as on a previously published experimental dataset that was annotated for spot locations. We have shown (i) that our approach achieves better performance when the training is performed on data with highly variable intensity and (ii) that performing training on a combination of experimental and simulated data is a viable approach in real-life setting.

Furthermore, we compared the performance of combining DeepSpot and ICY to that of the state-of-the-art deep learning-based method deepBlink and have shown that on average, DeepSpot enables a substantially better detection of mRNA spots than deepBlink. We found that DeepSpot / ICY workflow provided excellent quality spot detection on the test datasets corresponding to the datasets on which it has been trained, with the average F1-score above above 97%, but also achieved high precision results on fully unknown datasets with the F1-score of 88% for the datasets provided with the deepBlink publication. Taken together, the good results on both known and unknown data indicate that DeepSpot is a more generalist model than deepBlink and that it achieves a good balance between overfitting and underfitting. We hypothesize that this generalization capacity is possibly due to both strong regularization within the network and the diversity of signal provided by the carefully constructed training data.

To evaluate how well our method is suited for end-to-end biological investigations, we have shown the efficiency of the DeepSpot model trained on the combination of experimental and simulated data in the context of an independent study of cell migration. We have performed single molecule FISH to detect *β-Actin* in mouse fibroblasts in a wound healing assay and enhanced the resulting images using our combination model, which allowed us to detect that the *β-Actin* mRNA enrichment is specific to leading edge of migrating cells as contrasted by its expression in non-migrating cells.

To conclude, we have shown that DeepSpot enhancement enables automated detection and accurate localization of mRNA spots for downstream analysis methods and can thus be a useful tool to streamline not only spot detection, but also studies of localized mRNA enrichment within cells.

### Hardware and framework

All the analysis presented in this paper are performed on 1 GPU, Tesla T4 with 16GB of memory, on a dedicated machine with two 2 CPU Intel Xeon Silver 4114 and 128Go RAM. This work is implemented in Python 3.8 and we used TensorFlow 2.4 for creating and training the neural networks.

## Code and Data availability

DeepSpot network along with the code for training and for mRNA spot enhancement is fully open source and available on GitHub at https://github.com/cbib/DeepSpot. Our pretrained models used in this study are also available on our GitHub page.

Data for simulated images for DS_var_, DS_fixed_ and DS_hybrid_ are available on Xenodo at https://doi.org/10.5281/zenodo.5724466

## Acknowledgments

We thank Dr. Arthur Imbert for sharing the experimental smFISH data and helping to run the BIG-FISH pipeline.

